# Optimal Haplotype Assembly from High-Throughput Mate-Pair Reads

**DOI:** 10.1101/014993

**Authors:** Govinda M. Kamath, Eren Şaşoğlu, David Tse

## Abstract

Humans have 23 pairs of homologous chromosomes. The homologous pairs are almost identical pairs of chromosomes. For the most part, differences in homologous chromosome occur at certain documented positions called single nucleotide polymorphisms (SNPs). A haplotype of an individual is the pair of sequences of SNPs on the two homologous chromosomes. In this paper, we study the problem of inferring haplotypes of individuals from mate-pair reads of their genome. We give a simple formula for the coverage needed for haplotype assembly, under a generative model. The analysis here leverages connections of this problem with decoding convolutional codes.

## I. Introduction

Humans, like most mammals are diploid organisms, *i.e*. all somatic cells of humans contain two copies of the genetic material. Humans have 23 pairs of homologous chromosomes. Each member of a pair of homologous chromosomes, one paternal and the other maternal, has essentially the same genetic material, one exception being the sex-chromosomes in males. For the most part, homologous chromosomes differ in their bases (which take values in the alphabet {*A, C, G, T*}) at specific positions known as Single Nucleotide Polymorphisms (SNP). Other types of differences such as insertions and deletions are also possible, but these are rare occurrences, and are ignored in this paper. There are around 3 million known SNPs in humans, whose genome is of length approximately 3 billion base pairs. Thus on average a SNP appears once in 1000 base pairs. The positions where SNPs occur are well documented (see for instance [5]). Usually only one of two bases can be seen at any particular SNP position in the population. The one that is seen in the majority of a population is referred to as the *major allele*, and the other is referred to as the *minor allele*. The two sequences of SNPs, one on each of the homologous chromosomes, is called the *haplotype* of an individual. A person is said to be *homozygous* at position *i* if both homologous chromosomes have the same base at position *i*. Otherwise the person is said to be *heterozygous* at position *i*.

The haplotype of an individual provides important information in applications like personalized medicine, and understanding phylogenetic trees. The standard method to find a person’s haplotype is to first find her *genotype*, which is the set of allele pairs in her genome. Haplotype information, i.e., which allele lies in which chromosome, is then statistically inferred from a set of previously known haplotypes based on population genetics models [2]. Often, haplotypes of many individuals from a population are inferred jointly. This process is referred to as *haplotype phasing*. A major drawback of this approach is that many individuals in a population need to be sequenced to get reliable estimates of the haplotype of one person.

The advent of next generation sequencing technologies provides an affordable alternative to haplotype phasing. In particular, these technologies allow one to quickly and cheaply read the bases of hundreds of millions of short genome fragments, called *reads*. One can *align* such reads to a known human genome, thereby determining their physical location in the chromosome. Aligning a read to a reference does not reveal whether that read comes from the paternal or the maternal chromosome, however, hence the non-triviality of determining the haplotype.

Clearly, if a read covers zero or one SNP, it does not contain any information about how a particular SNP relates to other SNPs on the same chromosome, and is thus useless for haplotype assembly. That is, a read helps in determining the haplotype only if it covers at least two SNPs. This may seem like a problem at first sight, since as we mentioned above, adjacent SNPs are separated on average by 1, 000 bases, but most sequencing technologies produce reads of length only a few hundred bases. Fortunately, with somewhat more labour intensive library preparation techniques, one can produce *mate-pair reads*, i.e., reads that consist of two genome fragments separated by a number of bases. The length of the separation between the two segments of DNA read is known as the *insert size*. For popular technologies like Illumina, the read lengths are around 90 – 100 base pairs (bp), and the insert size ranges from around 300 bp to 10, 000 bp, with the median insert size around 3, 000 bp (See Figure 1, which was taken from [1]). These reads offer a possibility of inferring haplotypes from them. However, errors in reads present a major challenge. Both single and mate-pair reads are illustrated in Figure 2.

**Fig. 1.**
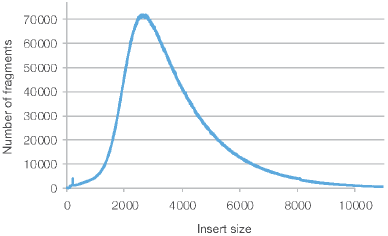
Insert size distribution of Human NA 12877 library sequenced on Illumina HiSeq® 2500. Median insert size was 3400 bp here. Figure taken from [1].

**Fig. 2.**
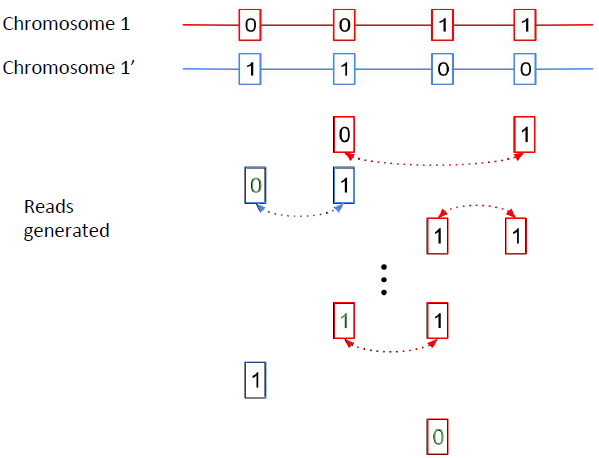
An illustration of read generation. The genetic material in chromosome 1 and chromosome 2 are identical apart from in the SNP positions shown. Here we represent the major allele by 0 and the minor allele by 1. Reads from chromosome 1 are represented in red and those from chromosome 2 are represented in red. We do not know which reads come from which chromosome. Mate-pair reads are indicated by dotted lines in between them. Errors are shown in green.

In this paper, we characterize the coverage, i.e., the number of reads that cover at least two SNP positions, required for haplotype assembly from paired reads. We determine this quantity as a function of the number of SNPs *n*, read error probability *p*, and the separation between the two reads in each mate-pair reads in terms of number of SNPs given by a random variable *W*. In particular, we show that for uniformly distributed read locations, the coverage required for a maximum-likelihood (ML) assembler to succeed is asymptotically

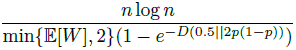

and no assembler can succeed on all possible haplotypes with fewer reads. In proving this result, we show the haplotype assembly problem’s connection to that of decoding a convolutional code, and use analysis methods for such codes. We note that we refer to natural logarithms by log throughout the manuscripts.

The haplotype assembly problem has been studied previously. In particular, that maximum-likelihood (ML) estimation can be done using dynamic programming in a similar setting was first recognized in [4]. The problem was also studied recently in [6]. There, it is assumed that a mate-pair could cover any two SNPs on the chromosome and bounds are derived on the number of reads needed to carry out haplotype assembly using a message passing algorithm. This however is an unrealistic assumption, as the chromosome lengths are several million, but the insert lengths are typically in the thousands. In this paper, we consider a model where reads only cover SNPs which are nearby.

## II. Noiseless reads covering adjacent SNPs

Consider first the case where all reads are noiseless. For convenience, we shall refer to the major allele by 1 and the minor allele by 0. We assume that the genome is heterozygous at all SNP positions, an assumption we will justify shortly. That is, if *S*_1_, *S*_2_, …, *S_n_* are the *n* SNPs on a chromosome, then the other chromosome has SNPs 
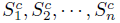
. Each mate-pair covers adjacent SNPs *i* and *i* + 1, where *i* is picked from {1,2, …, *n* − 1} uniformly at random. As mentioned above, the reads are assumed to be aligned to the reference genome, and therefore the genomic locations of all reads are known. A read covering SNP *i* and SNP *i* + 1, will output (*S_i_*, *S*_*i*+1_) with probability 
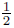
 and SNP 
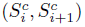
 with probability 
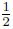
 depending on which of the two homologous chromosomes it is read from.

Determining the necessary and sufficient coverage in this setting is relatively simple. Indeed, note that the parity of *S_i_* and *S_i_*_+1_, which equals the parity of 
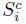
 and 
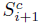
 is a sufficient statistic we get from reads covering SNP *i* and SNP *i* + 1. Thus we can reconstruct the haplotype when we see a read covering each of the *n* − 1 adjacent pair of SNPs. Note that this is equivalent to the coupon collector problem, and therefore reconstruction will be correct with high probability if one has *c*(*n* − 1) log(*n* − 1) reads with *c* > 1, whereas if *c* < 1, then reconstruction will fail with high probability.

## III. Noisy reads covering adjacent SNPs

Next, consider the case where each mate-pair again covers a pair of adjacent SNPs and aligned perfectly to a reference genome, but now each read in each pair is in error at the SNP position with probability *p* independent of everything else. Similarly to the noiseless case, we wish to characterize optimal coverage as a function of *p*. We will see that *O*(*n* log *n*) is the correct scaling here as well. In particular, we will say that *c*(*p*) is the optimal coverage if for any *∊* > 0, (*c*(*p*) – *∊*)*n* log *n* reads are insufficient to reconstruct with probability at least 1 – *∊*, but (*c*(*p*) + *∊*)*n* log *n* are sufficient as *n* → ∞. This makes optimal coverage only a function of *p*.

We again assume that each SNP position is heterozygous. This is a reasonable assumption since the positions of SNPs in humans are typically known, and thus one can test every SNP position for heterozygosity. This can be done using reads that cover a single SNP position, as we mentioned above, such reads are much easier to obtain in large numbers compared with mate-pairs. Thus, the coverage required for reliable heterozygosity testing can be met easily.

We will not fix the number of reads *M* to be *c*(*p*)(*n* − 1) ln *n* but instead allow *M* to be random, and in particular, be distributed as Poiss(*c*(*p*)(*n* − 1) ln *n*). This relaxation only simplifies the analysis and does not affect the result. Indeed, note that such a random variable is the sum of *n* ln *n* i.i.d. Poiss(*c*(*p*)) random variables, and therefore by the law of large numbers will take values in [*c*(*p*) ± *∊*] *n* ln *n* with high probability. We assume that SNP positions 1, 2, …, *n* − 1 are equally likely to be the first position covered by each read.

Here again, the set of parities of adjacent SNPs *i* and *i* + 1 are a sufficient statistic to reconstruct. As the probability of error in each SNP in each read is *p*, and errors are independent across reads, we have that the probability that a read gives us the wrong parity is

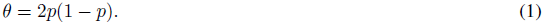

Let *L*_1_, *L*_2_, …, *L*_*n*−1_ denote the true parities, *i.e*. 
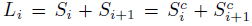
. For example, in Figure 2, *n* = 4, *S*_1_ = *S*_2_ = 0, *S*_3_ = *S*_4_ = 1, *L*_1_ *=* 0 = *L*_3_, *L*_2_ = 1. For *θ* ∈ [0,1], let the function *D*(*θ*) be defined as,

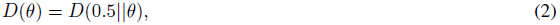

where *D*(·║·) denotes relative entropy measured in nats. Also let *Z*_*i*1_, …, *Z_i,N_i__* be the observations of *L_i_*, and note that due to Poisson splitting, we have *N_i_* ∼ Poiss(*c*(*p*) log *n*). Then we have that,

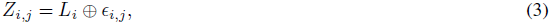

where *∊_i,j_* are Bern(*θ*) random variables independent of each other, and ⊕ denotes modulo-2 addition. The assembly problem therefore is equivalent to decoding the *L_i_*s from the observations *Z_i,j_*s, as in a repetition code, where each message symbol (*L_i_* 1 ≤ *i* ≤ *n* − 1) is repeated *N_i_* times.

Under these assumptions, we see that the maximum-likelihood (ML) rule is to declare *L_i_* = 0 (i.e., SNPs (*i*, *i* + 1) agree) if more than 
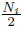
 of the observations are 0 and declare *L_i_* = 1 (SNPs (*i, i* + 1) disagree) otherwise. Note that each observation is correct with probability *θ*, and thus the number of correct observations ∼ Bin(*N_i_*, *θ*). Thus we have that,

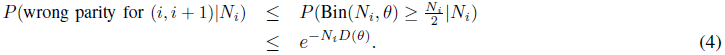

Assuming that 
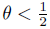
, we also have that (from chapter 12 of [3])

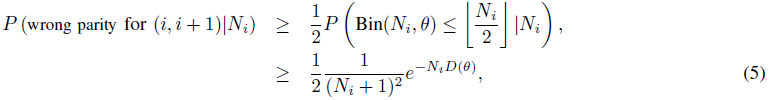

where we have used the fact that *D*(*p*║*q*) is a monotonically increasing of *p* for a fixed *q*, in the regime *p* > *q*. Some arithmetic then reveals that

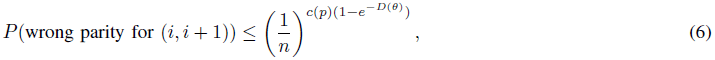

and that

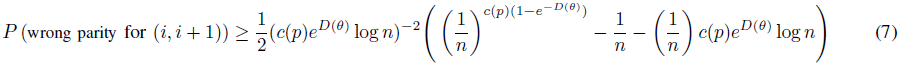

This leads to the following coverage result:

### Theorem 3.1

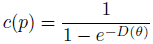

is the optimal coverage.

*Proof*: Note that

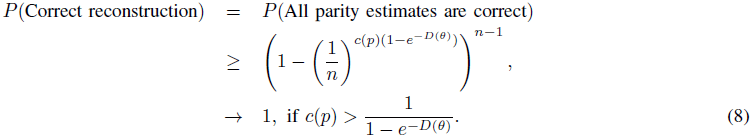

Further, we have

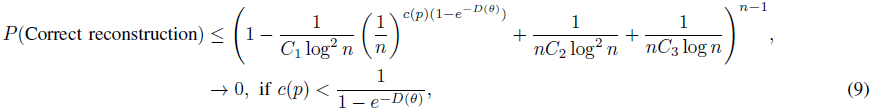

where *C*_1_, *C*_2_, *C*_3_ are positive constants as derived in (7).

## IV. Noisy reads covering non-adjacent SNPs

Next, we consider the case where mate-pairs cover more than just the adjacent SNPs. In particular, we will assume that each mate-pair covers *S_i_*, *S*_*i*+*W*_, where *i* is uniform as before, and *W* is a random integer between 1 and *w*, independently chosen for each read. *W* represents the separation between the two reads of a mate-pair read measured in terms of number of SNPs between them.

We will consider three cases, in order of increasing complexity:

### A. W is either 1 or 2 with equal probability

Let us first consider the case where we have observations of adjacent parities and skip-1 parities, *i.e*. parities *S_i_* + *S_i_*_+1_ and *S_i_* + *S_i_*_+2_ respectively. Let 
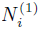
 be the number of noisy observations of *S_i_* + *S_i_*_+1_ and 
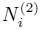
 be the number of noisy observations of *S_i_* + *S_i_*_+2_. Similarly to the previous case, each read consists of a uniformly chosen *i*, paired with *i* + 1 or *i* + 2 with probability 1/2. Therefore with a total of Poiss(*c*(*p*)*n* log *n*) reads, we have 
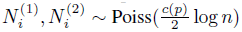
.

Let 
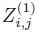
 be the *j*th noisy observation of *S_i_* + *S_i_*_+1_, and similarly 
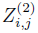
 be the *j*th noisy observation of *S_i_* + *S_i_*_+2_ (*i, i* + 2). That is,

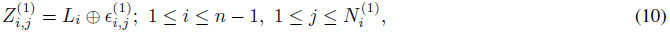

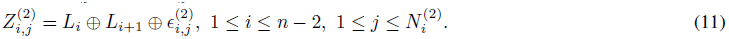

Note that the assembly problem is now equivalent to decoding the *L_i_*’s from noisy observations of the *L_i_*’s and (*L_i_* + *L_i_*_+1_)’s. Observe that this is equivalent to decoding a rate 
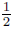
 convolutional code whose polynomial generator matrix is given by (Figure 3)

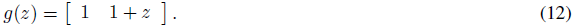

**Fig. 3.**
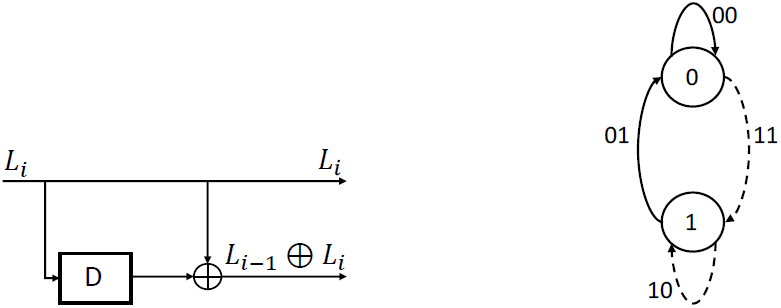
The convolutional code corresponding to the case where we see adjacent and skip-1 parities (on the left) and its state diagram (on the right). The transitions corresponding to input 0 is shown by solid lines and those corresponding to input 1 are shown by dashed lines. Each output here is seen correctly 
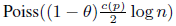
 times and incorrectly 
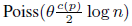
 times.

Each output bit of the code is repeated 
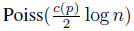
 times, and passed through a binary symmetric channel (BSC) with cross over probability *θ*. Unlike coding for communication, however, here one cannot initiate/terminate the code as desired, since the values of SNPs are given, and hence we can not 0 pad as in the communication setting. ML decoding can be done using the Viterbi algorithm, with a trellis that has 2 states (Figure 3). The points on each edge in the trellis corresponds to the number of observations which agree with the output of the transition between the states.

#### Theorem 4.1

In the setting above,

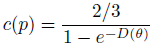

is the optimal coverage.

*Proof:* We will see that the error performance is determined by the *free distance* of the convolutional code, which is the minimum weight of any non-zero codeword. First note that this particular code is not catastrophic that is, no infinite-weight input produces a finite-weight output, as the GCD of the generator polynomials is 1. Thus to calculate the free distance, we can restrict ourselves to input message polynomials with finite degree. In particular, note that any input bits with weight more than 1 will have give rise to an output of weight at least 4. Any input with weight 1 gives us output of weight 3. Thus we conclude that the free distance of the code is 3.

Following the analysis in [7], consider the condition where the parties are *L*_1_ = *L*_2_ = … = *L*_*n*−1_ = 0. Next note that any state transition apart from the 0 → 0 transition adds an output of weight at least 1. Thus we note that the amount of weight accumulated by any path that diverged from the all 0 path ℓ stages ago accumulates weight at least *ℓ*.

Let *R_n_*(*p*) be the average number of reads that cover a position (that is an output of a convolutional code). We have that,

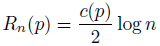

We further note that since the reads are all independently corrupted, the number of reads supporting the all zero path as compared with some weight *d* path depends only on the positions where the all zero path and the second path differ. We note that the reads supporting the weight *d* path at these points of difference from the all zero path is *X_d_* ∼ Poiss(*dθR_n_*(*p*)) (because the sum of independent Poisson random variables is a Poisson with rate equal to the sum of rates.). The points accumulated by the all zero path on these points is *X*_0_ ∼ Poiss(*d*(1 – *θ)R_n_*(*p*)). Further we have that *X*_0_╨*X_d_*, for all *d* > 1. Thus we have that the probability that a path of weight *d* is preferred over the all zero path with probability *P*(*X_d_* > *X*_0_), which can be bounded as (See Appendix A)

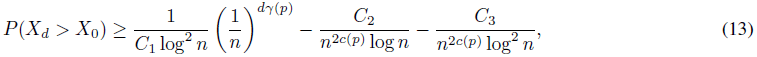

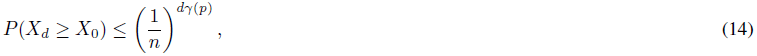

where 
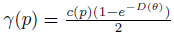
.

We first note that the all 0 path should not be killed by the weight 3 path that diverged from the all zero path 3 stages before the current stage. We note that there are 
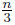
 disjoint events. Thus we have that,

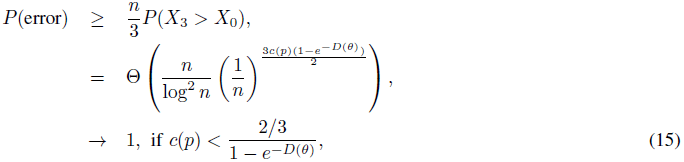

thus giving us that

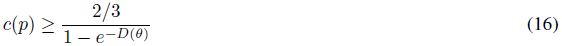

is necessary.

Next, we prove sufficiency. We say that the all-zero path is killed at stage *i* if some other path is preferred to the all 0 path at the 0 node in the trellis at stage *i*. At the last stage, this includes, the event that the path terminating at the 1 node of the trellis has accumulated more weight than the path terminating at the 0 node.

Let *p_i_* be the probability of the all-zero path being killed at stage *i* of the trellis. We will bound *p*_1_ + …, +*p_n_*.

First consider the case where the all zero path is killed in the first 3 stages. As there are only 8 paths in the trellis at this stage, the probability of this occurring is

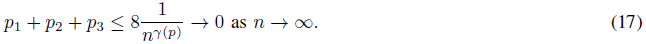

Next consider the probability of the all zero codeword being killed at the last stage. Note that each stage has at most 2 surviving paths, and any path of weight *w* has to have diverged from the all-zero path at most *w* stages ago, there are at most 2^*w*^ paths of weight *w* competing with the all zero codeword at any stage. Thus union bounding this probability is upper bounded as,

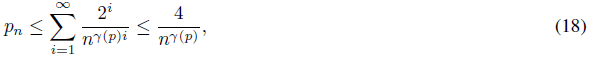

for large enough *n*, which also goes to 0 as *n* → ∞.

Finally consider the case of the all zero codeword being killed in the mid section of the trellis. Note that after the first 3 stages any path that did not diverge from the all zero path has hamming weight at least 3. Further note that from the definition of free distance, any path that diverged from the all zero path and is competing with the all zero path has weight at least 3. Thus the probability of the all zero codeword being killed at any stage is upper bounded by,

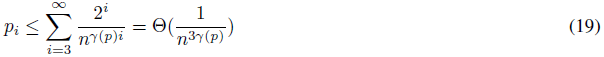

The total probability of error thus is asymptotically

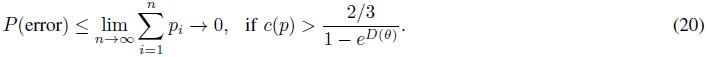

#### Remark 1

This result actually shows that asymptotically no algorithm can perfectly reconstruct even in this non-Bayesian setting when

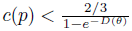
 for all assignments of *L*_1_, … *L*_*n*−1_.

To see this we first note that by the symmetry in the problem and the ML algorithm, the probability of error for every assignment of *L*_1_, … *L*_*n*−1_ is the same for the ML algorithm. Thus if any algorithm *A* succeeds to perfectly reconstruct when 
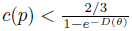
, then the probability of error of this algorithm on every assignment of *L*_1_, … *L*_*n*−1_, would have to be asymptotically lower than the probability of error of the ML algorithm on that assignment.

Further, we note that if each *L_i_* ∼Bern(0.5), then the ML algorithm would be the MAP algorithm, with the minimum probability of error. However this leads to a contradiction, because the probability of error of *A* in this Bayesian setting would then be asymptotically lower than that of the MAP algorithm.

### B. *W is uniform over* 1,…, *w for w* ≥ 3

We can observe adjacent, skip–1, skip–2, …, skip–(*w* − 1), parities, each being equally likely. Here let 
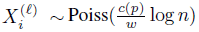
 be the number of observation of skip–*m* parities, (*i, i* + *m* + 1). With notation as before we have that, each observed parity can be represented as,

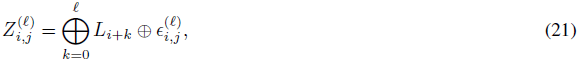

for 
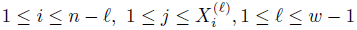
, where 
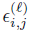
 are Bern(*θ*) random variables independent of each other.

We note that these can be represented by a rate 
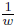
 convolutional code with polynomial generator matrix

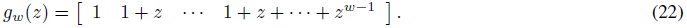

We further note that, the trellis corresponding to this code will have 2^*w*−1^ states.

#### Lemma 4.2

The free distance of the code whose polynomial generator matrix is given by *g_w_*(*z*) is *2w*, for *w* ≥ 3.

*Proof:* We note that, this code is not catastrophic, as the GCD of the polynomials is 1. Further, when *w* ≥ 3, we note that any monomial input will have output weight 
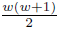
, which for *w* ≥ 3 is greater than 2*w*. Further, we note that for any input with more than 1 monomial, we will each term in the polynomial generator matrix will have outputs of weight at least 2. Thus we have that the free distance ≥ 2*w*. We note that the input 1 + *z* gives us an output of weight 2*w*. Hence, we have that the free distance of this code is 2*w*.

Following the exact same procedure as before, we have that,

#### Theorem 4.3

For the setting above, when *w* ≥ 3,

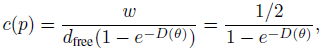

is the optimal coverage, where *d*_free_ is the free distance of the convolutional code, with polynomial generator matrix *g_w_*(*z*).

### C. W is non-uniform

Next we consider the case where, all parities are not observed in the same proportions. In particular, the separation between the two reads in a mate-pair measured in terms of number of SNPs between them is a random variable *W*, taking integral values between 1 and *w*, with probabilities *p*_1_,…, *p_w_*. That is, the number of observations of skip– (ℓ − 1) parities (*i, i* + *l*) is given by 
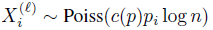
, where 
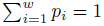
.

We further assume that the GCD of the generator polynomials corresponding to the support of *p*_1_, …, *p_w_* is 1. Let p = (*p*_1_, *p*_2_, …, *p_w_*).

To tackle this case we first for a general rate 
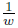
 convolutional code *C*, with *w* output streams, we define the averaged distance of two a codewords *v*_1_*, v*_2_ ∈ *C*, as

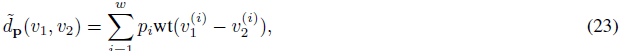

where 
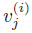
 is the codeword *v_j_* in the *i*-th stream, and 
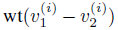
 is the hamming distance between *v*_1_ and *v*_2_ in the *i*-th stream. Further let *d̃*_*p*_(*v*) := *d̃p*(*v*, 0), be referred to as the averaged weight of a codeword *v* ∈ *C*

For any convolutional code *C*, define the averaged free distance,

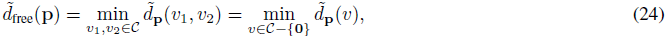

where the second equality follows from the linearity of *C*, and 0 is the all zero codeword. Henceforth, by abuse of notation, we shall represent *d̃*_free_(p) by *d̃*_free._

#### Lemma 4.4

If the GCD of the generator polynomials corresponding to the support of p = (*p*_1_, *p*_2_,…, *p_w_*) is 1 (*i.e.* the code is not catastrophic), for the family of codes under consideration (with polynomial generator matrix *g_w_*(*z*), *w* ≥ 2)), we have that,

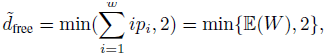

where 
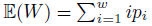
.

*Proof:* As the GCD of the polynomials corresponding to the support of p is 1, we have that the no input message polynomial of infinite weight can have an output codeword of finite averaged weight.

Further we note that for every input of more than 1 monomial the output on every stream will have at least weight 2, thus giving us that the averaged weight of any such message symbol is at least 2. Further, we note that the input monomial 1 + *z* will lead to a codeword with weight exactly 2 on each stream, giving us a codeword of averaged weight 2.

Next, we note that any input of 1 monomial will give rise a codeword of weight *i* on the *i*-th stream and hence gives a codeword with averaged weight 
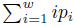
.

Thus we have that

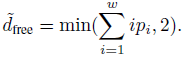

#### Theorem 4.5

In this case,

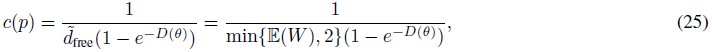

is the optimal coverage.

*Proof:* This is relegated to Appendix B.

#### Remark 2

On the ring of polynomials over the binary field 𝔽_2_[*z*], let,

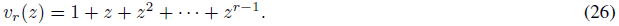

It is easy to see that if *r*|*s*, then *v_r_*(*z*)|*v_s_*(*z*). For 1 < *r* ≤ *s* ≤ 50, one can check that if *r* ∤ *s*, then *v_r_*(*z*) ∤ *v_s_*(*z*). We note that all polynomials in the polynomial generator matrix of the code considered here are of the form *v_r_*(*z*).

Thus we have that, for 1 ≤ *w* ≤ 50, the GCD of generator polynomials corresponding to the support of *p*_1_, …, *p_w_* is not equal to 1, only occurs when Support(*p*_1_, *p*_2_, …, *p_w_*) is a set of the form {1 < *j* ≤ *w* : *j* = *ki*, for some *k* ∈ ℕ}, for some integer *i* > 1. This corresponds to the case when our observations have a cyclic structure, in which case reconstruction is impossible.

## V. Non uniform SNP coverage

In all the above results, we have assumed that the number of reads covering every SNP position, with SNPs of its neighbourhood are identically distributed. In general, the genomic distance between adjacent SNP pairs will not be constant. There may be SNPs which do not have many SNPs in their near them on the genome, and hence may be covered by much fewer reads that cover multiple SNPs. We consider this in the setting of Section III, where only adjacent SNPs are covered by reads, as an illustration of how this can be handled. We first start with a simple example.

### Example 1

Suppose we only observe adjacent SNPs. If *α* fraction of adjacent SNPs were covered with probability 
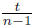
, *t* < 1 and the rest were covered with equal probability, then 
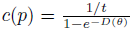
 is the optimal coverage. This can be shown using calculations identical to those of III.

Next, we consider a more general case. Suppose then that we only observe adjacent SNPs, and that the probability of each read covering SNPs (*i, i* + 1) is not 
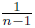
 but instead 
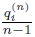
 (where 
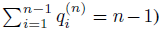
), and hence the number of observations of the *i*th parity is 
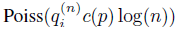
. Further assume that 
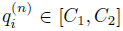
 for some constants 0 < *C*_1_ ≤ 1 ≤ *C*_2_. Fix *δ* > 0 and let 
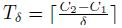
.

For any *δ* > 0, 1 ≤ *ℓ* < *T_δ_*, let,

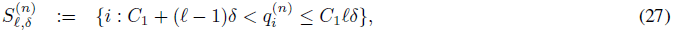

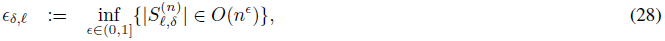

and define *m* and *k* as

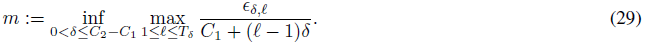

and

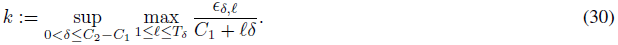

Then we have that

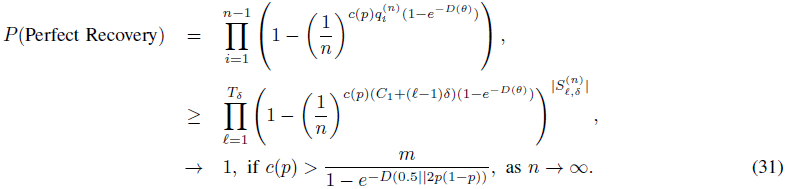

This gives us that

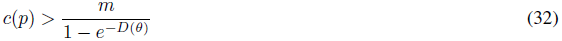

is sufficient for perfect reconstruction, and similar calculations to those of Section III give us that,

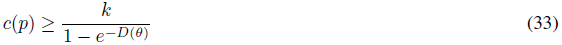

is necessary.

## Acknowledgements

This work is partially supported by the Center for Science of Information (CSoI), an NSF Science and Technology Center, under grant agreement CCF-0939370.

## Appendix

### A. Poisson Races

If *X* ∼ Poiss(λ) and *Y* ∼ Poiss(*µ*), *µ* > λ, *X* ╨ *Y* then from a Chernoff bound, we have that,

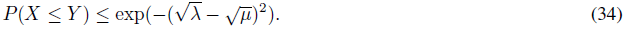

Further noting, that *X* + *Y* ∼ Poiss (λ + *µ*). And 
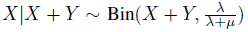
. One can show that,

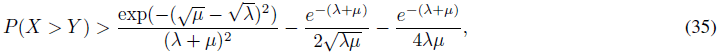

by noting that 
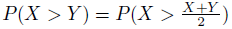
, and upper bounding that term conditioned on *X* + *Y* = *i*, ∀*i* ∈ ℤ.

### B. Proof of Theorem 4.5

As in proof of Theorem 4.1, the probability that a averaged weight *d* path is preferred to the all zero path is given by (as this is a race between a the number of erroneous reads at positions of difference ∼ Poiss(*dθc*(*p*) ln *n*) and the number of correct reads at positions of difference ∼ Poiss(*d*(1 – *θ*)*c*(*p*) ln *n*).),

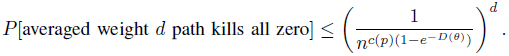

Let,

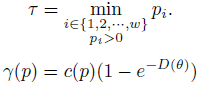

We thus have that in every 2^*w*−1^ + 1 stages any path on the trellis that does not visit the all zero stage adds a averaged weight of at least *τ* (because code is not catastrophic implies, that we can not get from any state other than the all zero state to itself, without adding any weight, and one has to visit some state twice in 2^*w*−1^ + 1 stages). Let 
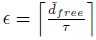
. Thus the number of paths in the trellis that have averaged weight less than *d̃*_free_ + ℓ*τ*, is at most 2^*w*(2^*w*−1^+1)(ℓ + *∊*)^.

As in the proof of Theorem 4.1, we can show that the probability that the all 0 path will be killed in the first 2^*w*(2^*w*−1^+1)*∊*^ stages or the last stage go to 0 as *n* → ∞. We now restrict our attention to the all 0 path being killed in the middle stages, where any codeword competing with the all 0 codeword has averaged weight at least *d̃*_free_.

The probability of a averaged weight *d* killing the all zero path is less than the probability that a averaged weight *d̃*_free_+ ℓ*τ* path kills the all zero, where ℓ is picked such that *d̃*_free_ + ℓ*τ* < *d* ≤ *d̃*_free_ + (ℓ + 1)*τ*. Thus we have that, the probability of error at any stage here is,

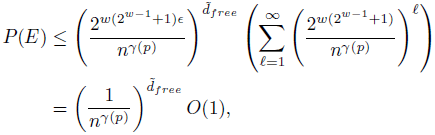

for large enough *n*.

Thus, by union bounding the probability of error across all stages as before, we can show that if

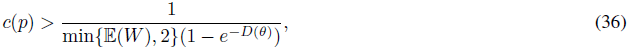

then we have perfect reconstruction.

The argument for necessity is essentially the same as that of Theorem 4.1, with a similar quantization argument.

